# Microglia maintain retinal redox homeostasis following ablation of rod photoreceptors in zebrafish

**DOI:** 10.1101/2025.10.17.683173

**Authors:** Michael Morales, Diana M Mitchell

## Abstract

Microglia rapidly respond to injury, stress, and perturbations to neurons in the brain and retina and perform phagocytosis to clear dying cells and debris. Oxidative stress is a frequent feature of neurodegeneration, and while glia are crucial for managing such stress, microglia may also be dysfunctional in diseased tissue. Here we examine the role of microglia in management of oxidative stress upon death of rod photoreceptors in the larval zebrafish retina. Using *rho*:nfsb-eGFP transgenic zebrafish and treatment with the pro-drug metronidazole (MTZ), we coupled the generation of reactive oxygen species (ROS) in dying rods to their ablation. Microglia efficiently engulfed and cleared the ROS-laden rods, effectively undertaking the oxidative load. Despite abundant ROS upon MTZ-mediated cell death, oxidative stress overall was minimal in retinal tissue when microglia were present, indicating that they rapidly and efficiently performed redox functions. In *irf8*-/- mutants, which are deficient in microglia, retinas with MTZ-induced rod ablation showed widespread ROS that localized, at least in part, to Müller glia. Further, there was evidence of increased oxidative stress, and increased numbers of “off-target” inner retinal neurons that stained positive for the cell death marker TUNEL. Supplementation with the antioxidant Glutathione (GSH) reduced the number of off-target TUNEL+ cells detected in microglia-deficient retinas following rod ablation. Our results indicate that microglial redox functions are important in restoring homeostasis following acute retinal damage.

## Introduction

Microglia are professional phagocytes residing in the central nervous system of vertebrates. They play an important role in brain and retinal development through clearance of apoptotic and non-apoptotic neurons [1–6] and by shaping neuronal synapses [7–9]. Microglia respond to neuronal damage resulting from injury and disease by migrating to the site of lesion, clearing dead cells and debris through phagocytosis, and secreting cytokines [10–13]. Microglia limit secondary cell death following blunt impact trauma to the zebrafish brain [14] and limit death of photoreceptors following retinal detachment [15], findings that underscore their significance in restoring homeostasis following acute injury. Further, microglia are a prominent area of interest in neurodegenerative disease, where their functions may be beneficial or detrimental to pathological outcomes [16]. For example, microglia may coordinate cellular interactions in tissue repair [17] or in contrast they may show dysregulated phagocytic activity [18].

Both acute tissue damage and chronic neurodegeneration are associated with increased reactive oxygen species (ROS) and oxidative stress [19–22]. Dying or damaged cells may be limited in their ability to execute antioxidant mechanisms, thus other cell types present in the tissue must perform antioxidant functions to limit pathology [23]. It is also thought that neurons have a limited capacity to manage ROS [24], and there is evidence that neurons may transfer oxidized material to glia to manage oxidative stress[25–29]. Further, glia supply neurons with antioxidants [30]. Given the phagocytic and scavenging activity of microglia, microglia are often directly challenged with ROS and must appropriately manage oxidative stress.

We previously published a transcriptome analysis of microglia isolated from zebrafish retinas following neuronal death induced by the neurotoxin ouabain [31]. Among the genes expressed by microglia in this context were those with antioxidant function, such as glutathione peroxidase 1a (*gpx1a*), glutathione reductase (*gsr*), xCT (*slc7a11*), heme oxidase 1a (*hmox1a*), p62 (*sqstm1*), and the transcription factor nuclear factor erythroid 2-like 2 (*nfe2l2*), commonly known as *Nrf2* which regulates the expression of many antioxidant genes [32–36]. The expression of such transcripts by microglia indicates an antioxidant function and possibly an important role of microglia in controlling oxidants such as ROS in damaged retinas.

Microglial responses in the zebrafish retina have been studied using genetic mutants that are deficient in microglia [37–39] as well as pharmacological manipulation of microglia [37,40,41]. These studies primarily focused on microglia in regenerative responses of the zebrafish retina, with less emphasis on early microglial functions in response to neuronal death. Given the knowledge gap in this area, as well as the antioxidant gene expression noted previously, we therefore examined the role of retinal microglia in response to photoreceptor death that involves the generation of ROS. Our experiments were facilitated using a transgenic zebrafish line *rho*:nfsb-eGFP*^nt19* [42], in which rod photoreceptors (rods) in the retina express the nitroreductase enzyme nfsB. The reaction of the pro-drug metronidazole (MTZ) with nitroreductase enzymes is thought to produce ROS [43], and another study using zebrafish demonstrated ROS generation in nfsB-expressing β-cells of the pancreas when exposed to MTZ [44].

We hypothesized that microglial antioxidant function facilitates the recovery of retinal tissue following the ROS-generating death of retinal neurons. To investigate the function of microglia in this context, we used *rho*:nfsb-eGFP transgenic, *irf8^st95* homozygous mutant larval zebrafish [45,46], which are deficient in retinal microglia [46]. We demonstrate that MTZ mediated death of rod photoreceptors is ROS generating, and that microglia increase their oxidative load in response to rod ablation, which is coupled to microglial engulfment of dying rods. In *irf8* mutants that lack microglia, we found that ROS became widespread upon rod ablation, and this was associated with the detection of increased numbers of dying neurons in the retina, many of which were “off-target” HuC/D+ inner retinal neurons that did not express the *rho*:nfsB transgene. Supplementation with glutathione (GSH) in *irf8* mutants following MTZ-mediated rod ablation reduced the number of “off-target” retinal neurons detected following rod ablation, indicating that the lack of antioxidant function was at least in part responsible for the consequences observed in microglia deficient mutants. Collectively, our results indicate that antioxidant functions of microglia are essential to the recovery of retinal tissue following neuronal damage.

## Results

### Rod ablation upon metronidazole exposure of *rho*:nfsb-eGFP larvae generates an oxidative load in microglia

To induce the death of rod photoreceptors, we immersed *rho*:nfsb-eGFP*^nt19* [42] larval zebrafish at 3.5 days post fertilization (dpf) in a 10mM metronidazole (MTZ), 0.1% DMSO solution prepared in fish system water. Death of rod photoreceptors by 48 hours post-treatment (hpt) with MTZ was readily visualized by TUNEL staining (Figure 1A-B). The metabolism of MTZ by the nfsB enzyme has previously been shown to generate ROS in other cell types [44], and microglia are known to engulf dying retinal neurons [1,2]. To visualize reactive oxygen species (ROS) and microglia upon rod ablation in larval zebrafish retinas, we used CellRox™ Deep Red, an ROS probe that has been reported to detect superoxide and hydroxyl radicals [47,48] followed by immunostaining with the 4C4 antibody. Following exposure to MTZ or DMSO, then staining, whole larval *rho*:nfsb-eGFP eyes were dissected and imaged by confocal microscopy (Figure 1C-F’’’). Consistent with generation of ROS by the nfsB/MTZ reaction, CellRox™ signal was not observed in rods following DMSO treatment (Figure 1C, C’) but was readily detected in collapsed GFP+ rods following MTZ treatment (Figure 1D, D’). In DMSO treated eyes, microglia were located throughout the retina, and in some microglia, CellRox™ fluorescence was detectable in small, discrete puncta (Figure 1E-E’’’), suggesting that microglia generate or scavenge ROS-generating oxidants in developing retinas. In MTZ-treated eyes, microglia localized primarily to the ventral retina which also showed strong CellRox™ signal; this signal was often co-localized to GFP+ collapsed rods and often putatively within microglial compartments (Figure 1F-F’’’). The ventral region of the developing zebrafish retina is rich in rod photoreceptors [49,50] (visible in Fig 1E), and these images indicate that microglia are engulfing dying rods within this region following MTZ exposure. We measured the CellRox™ fluorescence intensity that was co-localized to microglia and found that the microglia in MTZ-treated eyes have increased CellRox™ fluorescence intensity compared to microglia in DMSO-treated eyes (Figure 1G). These results indicate that microglia responding to MTZ-induced rod cell death are challenged with an increased oxidative load, likely due to engulfment of ROS-laden rod photoreceptors.

**Figure 1:**
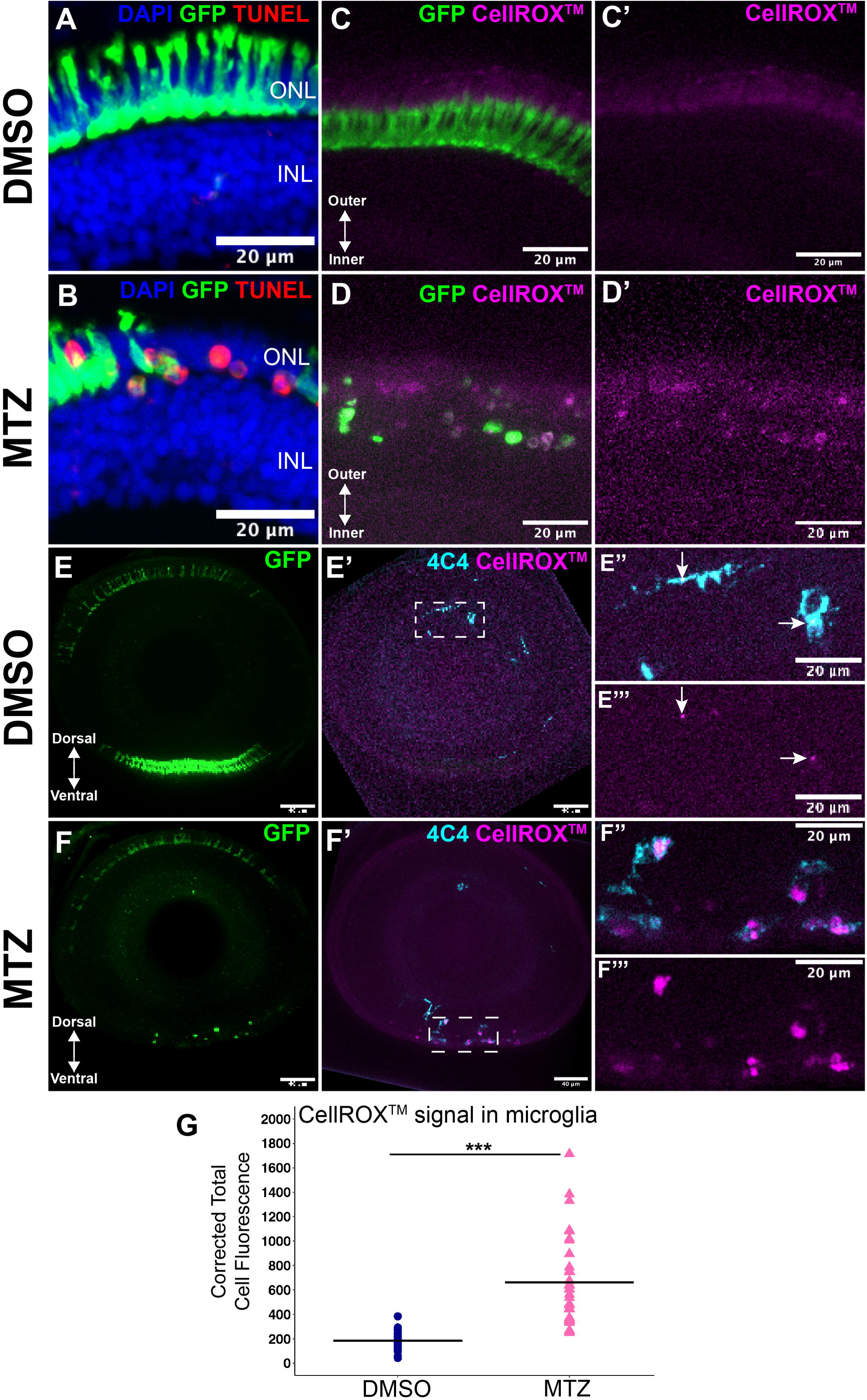
Rod ablation upon MTZ exposure of *rho:nfsb-eGFP* larval zebrafish increases oxidative load in microglia. A-B. Selected regions of retinal cryosections from DMSO (vehicle) treated controls (A) and following MTZ-treatment at 48 hours post treatment, hpt (B). In MTZ treated samples, rod cells (GFP+) are collapsed and co-localize with TUNEL labeling (red). DAPI stain shows cell nuclei. ONL: outer nuclear layer, INL: inner nuclear layer. **C-C’:** Region of ventral patch rods in a whole eye following DMSO treatment at 48 hpt showing rod cells (green), and CellRox™ (magenta). **C-C’:** Region of rods and CellRox™ signal in ventral retina from a whole eye confocal images following DMSO treatment at 48 hpt. **D-D’:** Region of rods in ventral retina from a whole eye confocal images following MTZ treatment at 48 hpt. Rods (green) often appear collapsed and co-localize with CellRox™ signal. **E:** Whole eye image showing GFP+ rods following DMSO treatment at 48 hpt. **E’:** The same image but showing channels representing microglia (4C4 stain, cyan) and CellRox™ signal (magenta). **E’’-E’’’**: Selected enlargement from inset region indicated in E’. Separate channels to visualize CellRox™ signal as puncta inside of microglia (arrows). **F:** Whole eye image showning GFP+ rods (green) following MTZ treatment at 48 hpt. **F’:** The same image but showing channels representing microglia (4C4 stain, cyan) and CellRox™ signal (magenta). **F’’-F’’’:** Selected enlargement from inset region indicated in F’. Separate channels to visualize CellRox™ signal associated with microglia, localized to putative microglial compartments. **G:** Fluorescence intensity measurement of CellRox™ signal inside microglia cells measured from DMSO treated or MTZ treated samples. Each datapoint represents a fluorescence measurement from a single microglial cell; microglia were selected from 4-5 of retinas in each treatment group. ***p<0.001, student’s t-test.

Oxidative stress is managed through a cellular response involving specialized enzymes that execute oxidation-reduction (redox) reactions to neutralize ROS and free radicals [32,51]. We therefore used hybridization chain reaction fluorescent in-situ hybridization (HCR-FISH) with probe sets to detect mRNA for *glutathione peroxidase 1a (gpx1a)* and *glutathione reductase (gsr),* two enzymes that perform redox reactions oxidizing sulfhydryl glutathione (GSH) to a disulfide dimer (GSSG) in order to reduce ROS, and the NADPH-dependent reduction reaction to restore GSH [52], respectively. To label microglia in these samples, we used probe sets to detect *mpeg1*, a gene known to be expressed in zebrafish microglia [1,2,13,31,46,53–55]. In DMSO-treated samples, signal from both *gpx1a* and *gsr* were detected within regions also co-labeling with *mpeg1* (Figure 2A, B, arrows), indicating that microglia express these genes in developing retinas. In MTZ-treated eyes, signal for *gpx1a* and *gsr* appeared increased within microglia localized to the rod-dense ventral patch (Figure 2A’, B’, arrows). We quantified the fluorescence intensity of *gsr* and *gpx1a* signal within regions also occupied by *mpeg1* signal and found that signal for both *gsr* and *gpx1a* were increased in *mpeg1*+ cells in MTZ-treated retinas compared to DMSO-treated samples (Figure 2E, 2F). These results further indicate that microglia respond to increased oxidative load resulting from widespread rod cell death induced by MTZ treatment.

**Figure 2:**
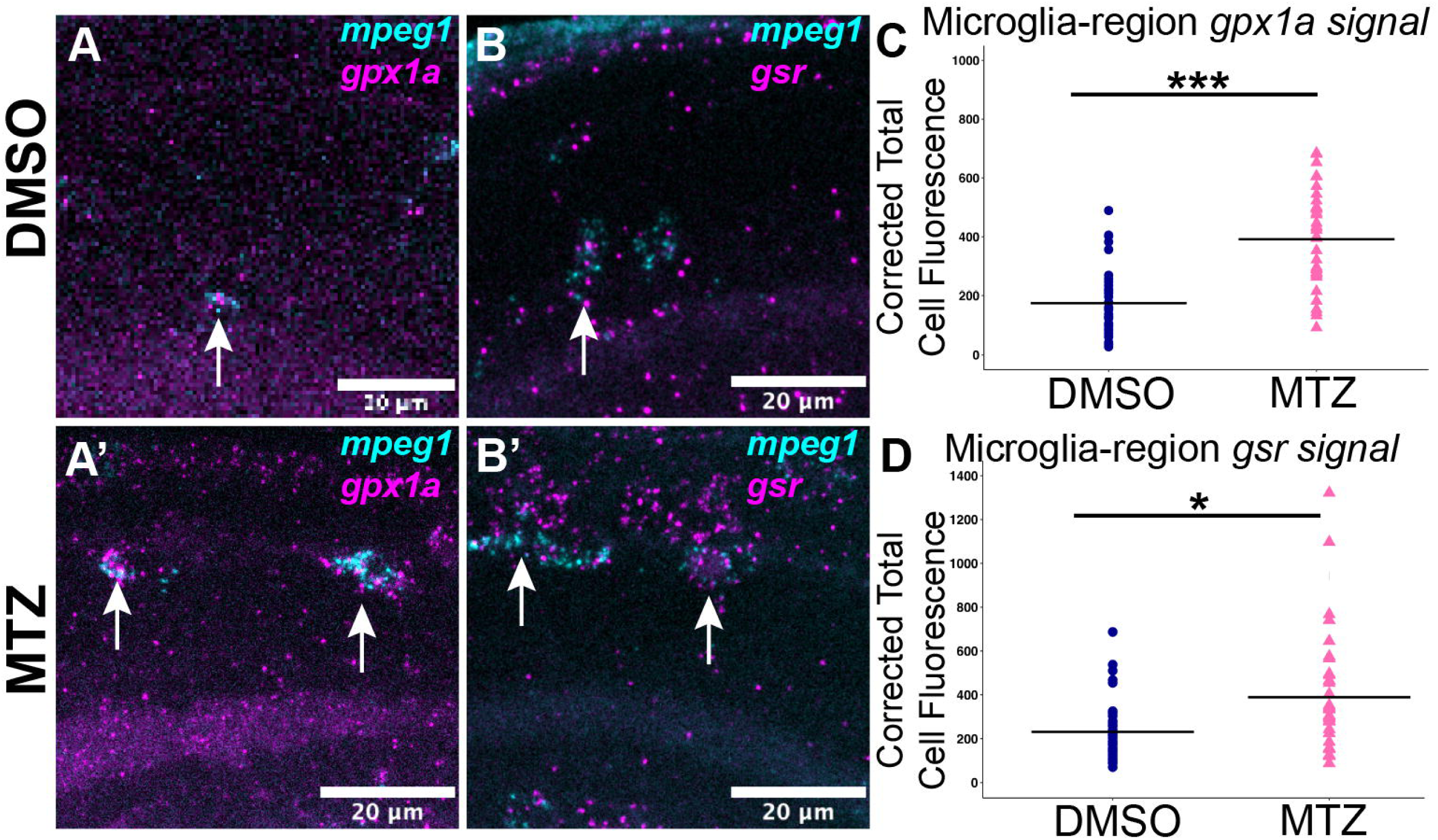
Detection of mRNA for antioxidant genes *gpx1a* and *gsr* in microglia increases following MTZ-mediated rod ablation. A-B: Images from confocal imaging of whole larval eyes following RNA fluorescence in situ hybridization (FISH) to label mRNA transcripts for *gpx1a* (magenta, A) and *gsr* (magenta, B) and *mpeg1* (microglia, cyan) in DMSO-treated (control) retinas at 48 hpt. **A’-B’:** Images from confocal imaging of whole larval eyes following RNA-FISH to label mRNA transcripts for *gpx1a* (magenta, A’), *gsr* (magenta, B’), and *mpeg1* (microglia, cyan) in MTZ-treated retinas following rod ablation at 48 hpt. **C-D:** Corrected total cell fluorescence intensity quantifications of signal from *gpx1a* (C) and *gsr* (D) mRNA transcripts co-localized with regions of *mpeg1* transcripts in DMSO and MTZ-treated retinas, with transcripts from MTZ samples measured primarily in the ventral patch. Each datapoint represents a fluorescence measurement from a single microglial region determined by *mpeg1* transcript signal; microglia were selected from 4 retinas in each treatment group.*p<0.05, ***p<0.001, student’s t-test.

### Reactive oxygen species are widespread in microglia-deficient retinas following rod ablation

We used microglia deficient *irf8* homozygous mutant (*irf8*-/-) zebrafish [45,46] to determine the consequences of microglia deficiency following the ablation of rods. Through genetic crosses, we generated clutches of larvae carrying the *rho*:nfsb-eGFP transgene that were either *irf8*+/-(microglia-sufficient) or *irf8*-/- (microglia-deficient). Clutches were split into two groups and treated with DMSO or MTZ for 48 hours, with samples collected at 24 and 48 hpt, and genotyped for *irf8* mutation post-collection. We again used CellRox™ and 4C4 to label ROS and microglia, respectively, in whole larval eyes which were then imaged by confocal microscopy. Consistent with results described in Figure 1, CellRox™ signal was not significantly detected following DMSO treatment of *irf8*+/- larvae (Figure 3A, A’). Microglia-sufficient (*irf8*+/-) eyes treated with MTZ showed increased CellRox™ signal in regions with dying rods, particularly in ventral retina where rods are densely populated, and microglia were observed to localize to the ventral region (Figure 3B-C’). These changes were detected by 24 hpt and remained visible through 48 hpt (Figure 3B, B’,C,C’). Rods were clearly detectable in *irf8*-/-eyes following DMSO treatment, while 4C4+ microglia were not present (Figure 3D). In *irf8*-/- eyes treated with MTZ, CellRox™ signal was concentrated heavily in the ventral region where dying rods were localized (Figure 3E-F’), but also readily detected throughout the eye/retina (Figure 3E-F). This widespread CellRox™ signal in *irf8*-/- eyes was detectable by 24 hours of MTZ treatment and became more pronounced at 48 hpt (Figure 3E-F). Also notable in *irf8*-/- eyes was the near absence of 4C4 staining in both DMSO and MTZ samples (Figure 3D-F), confirming microglia-deficiency in this mutant and the absence of additional 4C4+ cells that might respond to rod death.

**Figure 3:**
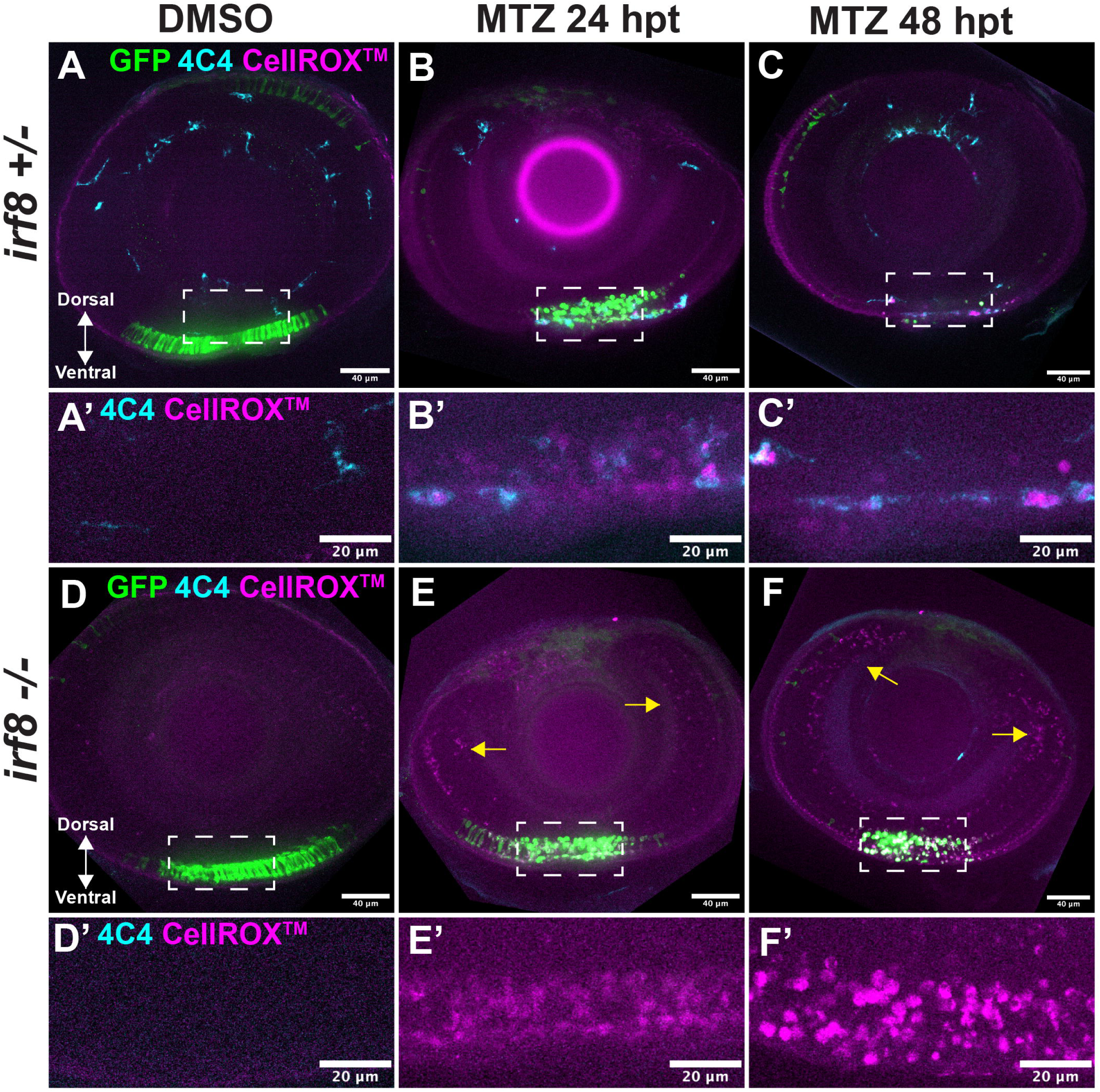
ROS are widespread in larval zebrafish retinas following Mtz-mediated ablation of rods in microglia-deficient conditions. A-C’: Rods, CellRox™ and microglia (4C4) visualization in whole eyes imaged by confocal microscopy, from *rho:nfsb-eGFP;irf8+/-* larvae following DMSO (A) and MTZ-treatment (B-C), at 24 and 48 hpt, respectively. DMSO images are shown for 24 hpt. A’-C’ represent enlarged regions corresponding to selected insets in A-C, showing the CellRox™ and 4C4 signal in the ventral retina. D-F: Rods, CellRox™ and microglia (4C4) ) visualization in whole eyes imaged by confocal microscopy, from *rho:nfsb-eGFP;irf8-/-* larvae following DMSO (D) and MTZ-treatment (E-F), at 24 and 48 hpt, respectively. DMSO images are shown for 24 hpt. Yellow arrows indicate CellRox™ signal in the inner retina observed in *irf8-/-* retinas. D’-F’ enlarged regions corresponding to selected insets in D-E, showing the CellRox™ and 4C4 signal in the ventral regions.

To determine if the ROS signal in *irf8* mutants was caused by metronidazole exposure alone (and not dependent upon activity of the nfsB enzyme), we exposed non-transgenic *irf8-/-* fish to the same MTZ concentration as used for rod ablation, for 48 hours. In the absence of nfsB expression, microglia deficient eyes lacked CellRox™ signal in the ventral patch region (Supplemental Figure 1), confirming that ROS generation in the ventral patch is a result of the MTZ/nfsB reaction. CellRox™ signal was detected in other regions of the eye/retina, but it was visibly weaker than what was seen for MTZ treated *rho*:nfsb-eGFP*;irf8-/-* eyes (Supplemental Figure 1). Overall, these results indicate that the absence of microglia leads to widespread ROS throughout the retina following MTZ-mediated ablation of nfsB-expressing rods, indicating that microglia are required to regulate ROS in this context. We further used samples of *rho*:nfsb-eGFP;*irf8-/-* larvae exposed to MTZ to determine if the CellRox™ signal was localized to Müller glia, because Müller glia may compensate for microglial function in their absence [46]. Use of an antibody for glutamine synthetase (GS), a Müller glia cell marker, revealed that the widespread ROS signal in *irf8*-/- eyes showed many instances of co-localization with Müller glia cell bodies and processes (Movie 1). Further, visualization of z-stack optical sections from whole eyes including the nuclear stain DAPI revealed that ROS signal in *irf8*-/-, but not *irf8*+/-, eyes also often co-localized with apparent pyknotic nuclei that were abundant in the inner retinal layers of *irf8*-/- eyes (Movie 2).

Collectively, these results indicate that dying rods are a source of ROS following MTZ-mediated ablation, that microglia control ROS upon rod ablation, and that clearance of dying cells and/or potentially cell viability are impacted by the absence of microglia.

### Increased numbers of off-target TUNEL+ cells are detected in microglia-deficient *rho*:nfsb-eGFP retinas treated with MTZ

To further investigate the impacts of microglia deficiency following MTZ-mediated ablation of rods, and given the presence of pyknotic nuclei described above, we used the TUNEL assay on retinal cryosections collected from *rho*:nfsb-eGFP fish following 48 hours of DMSO or MTZ exposure (Figure 4). We quantified the number of TUNEL+ cells in entire retinal cryosections using samples from the two *irf8* genotypes following DMSO or MTZ treatment. In DMSO treated samples, we found that *irf8-/-* retinas had increased numbers of TUNEL+ cells overall compared to *irf8+/-* retinas (Figure 4D), consistent with the loss of clearance function of retinal microglia [2,46]. Following rod ablation by MTZ treatment, the number of TUNEL+ cells increased in both genotypes, but this was most significant in *irf8*-/- retinas (Figure 4A-C’, 4D). We therefore examined TUNEL signal within the three defined retinal layers: outer nuclear layer, inner nuclear layer, and ganglion cell layer (ONL, INL, and GCL, respectively). In *irf8*+/- retinas, an increase in TUNEL+ cell numbers was detected only in the ONL (Figure 4E), consistent with on-target ablation of rods. The number of TUNEL+ cells in the ONL of *irf8*-/- retinas was further increased compared to *irf8*+/- (Figure 4E), which would be expected in the absence of microglial clearance of targeted rod photoreceptors. However, in *irf8*-/- retinas, TUNEL+ cells were also detected in the INL, and in the GCL, near and within the ciliary marginal zone (CMZ, Figure 4A’, B’, F-H). We noted that the TUNEL+ cells appeared to cluster, primarily in inner retinal regions in the periphery and near the CMZ region (Figure 4A-C’). Quantifications confirmed the presence of increased numbers of TUNEL+ cells in the INL and GCL of *irf8*-/- retinas following rod ablation/MTZ treatment, which was not detected in *irf8+/-* retinas (Figure 4F, G, H).

**Figure 4:**
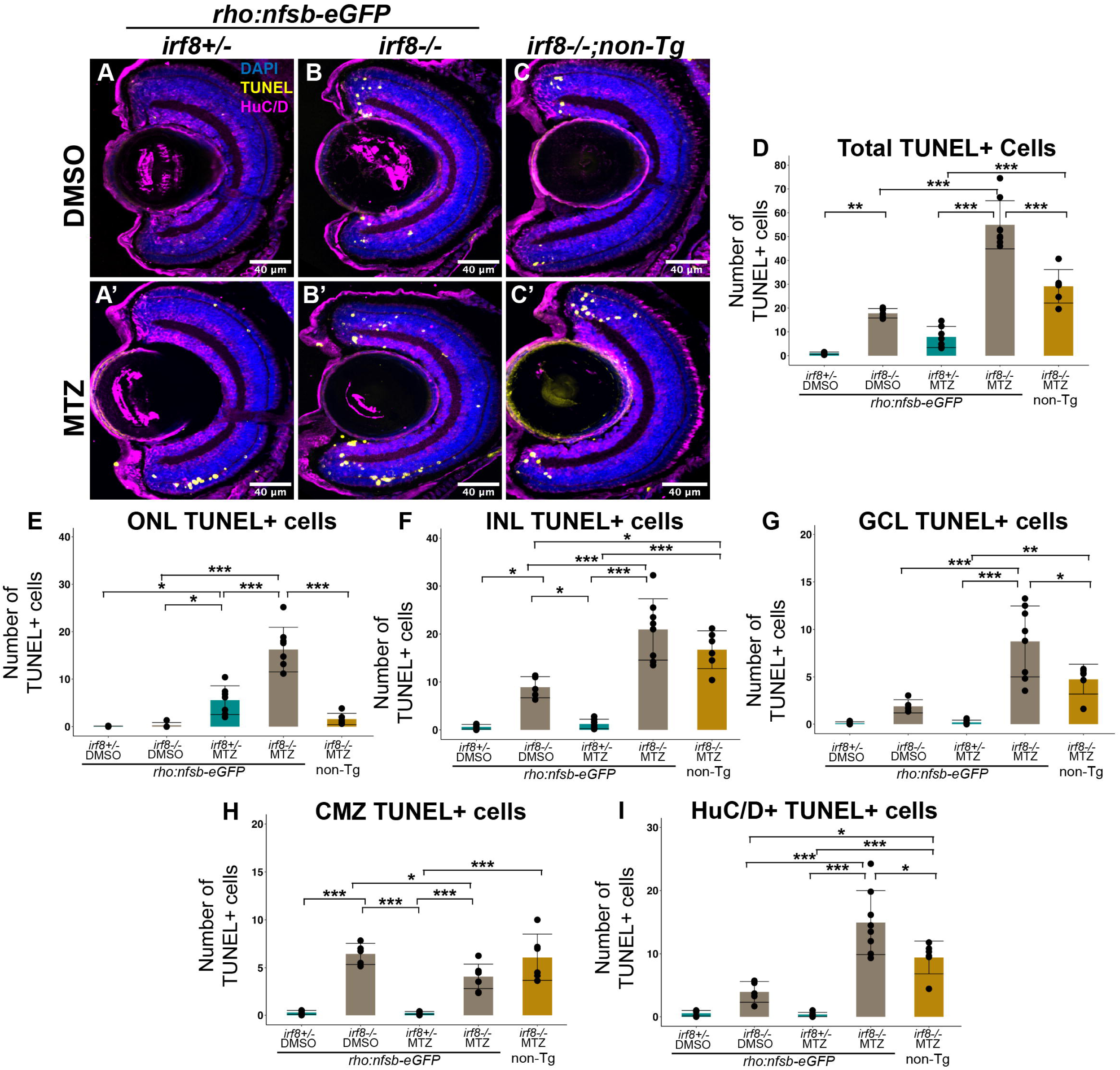
Cell death marker TUNEL shows increased detection outside of the outer nuclear layer in microglia deficient *rho*:nfsb-eGFP retinas. A-C’: Representative images of retinal cryosections stained for TUNEL (yellow), DAPI (blue), and HuC/D (magenta) from DMSO-treated (A-C) or MTZ-treated (A’-C’) larvae belonging to either *rho:nfsb-eGFP;irf8+/-* (A, A’), *rho:nfsb-eGFP;irf8-/-* (B, B’), or non-transgenic (non-Tg); *irf8-/-* (C, C’) genotypes. D-I: Quantifications of the number of TUNEL+ cells per retinal cryosection by indicated region at 48 hpt. Bars represent mean +/- SD, and dots represent counts from individual eyes. *p<0.05, **p<0.01, ***p<0.001; two-way ANOVA with Tukey’s HSD post-hoc test for multiple comparisons. For clarity, only statistically significant differences are annotated in these graphs.

To better identify TUNEL+ cells in the inner retina, we stained for HuC/D, a marker for amacrine and ganglion cells [56] (Figure 4A-C’, magenta). These images revealed that many of the TUNEL+ cells in the inner retina of *rho*:nfsb-eGFP;*irf8*-/- fish treated with MTZ are HuC/D+ neurons (Figure 4I), indicating that microglia deficiency increases the number of off-target TUNEL+ neurons detected following rod ablation. To again determine if this was related to MTZ-treatment alone, we included an analysis of non-transgenic (non-Tg, not expressing nfsB) *irf8-/-* fish treated with MTZ. The levels of TUNEL+ cells were not significantly different from the *irf8*-/- DMSO controls for total TUNEL+ cells and TUNEL+ cells in the ONL but were increased within the INL and specifically for HuC/D+ TUNEL+ neurons, though the levels were less than *irf8*-/- with MTZ-mediated rod ablation (Figure 4A-I). Collectively, these results indicate that upon metronidazole exposure and the death of rods, microglia deficiency results in increased numbers of TUNEL+ cells in retinal layers outside of the ONL which represent regions outside of the location of nfsB-expressing rods (“off-target” cell death detection).

To determine when increased TUNEL+ cells become detectable in microglia-deficient retinas, and if such numbers are reduced or resolved over time, we next performed a similar analysis of TUNEL+ cell numbers at 24-hour intervals from 24-72 hpt following DMSO or MTZ treatment of *rho*:nfsb-eGFP retinas of either *irf8+/-* or *irf8-/-* genotype (Figure 5). In *irf8*+/- retinas, increased TUNEL+ cells were detected at 24 and 48 hpt following MTZ treatment, primarily localized to the ONL with decreasing numbers through 72 hpt, consistent with rod death by 24 hpt and subsequent clearance (Figure 5A, B). By 24 hours of MTZ treatment, there were increased overall numbers of TUNEL+ cells in *irf8-/-* retinas relative to DMSO controls of both genotypes and compared to *irf8*+/- treated with MTZ, and these numbers remained elevated through 48 hpt (Figure 5A). Examining the trajectory of TUNEL+ counts in the ONL indicated that clearance of dying rods from MTZ treatment was delayed in *irf8*-/- mutants compared to *irf8*+/- but still obtained by 72 hpt (Figure 5B). In the INL and GCL, the TUNEL+ counts were elevated in *irf8-/-* mutants in both DMSO and MTZ groups compared to *irf8*+/- at 24 hpt, with only the elevation in TUNEL+ cells observed in the INL in the MTZ-treated group being statistically significant (Figure 5C,D). At 48 hpt, TUNEL+ cells remained elevated overall (Figure 5A), in the INL (Figure 5C), and in the GCL (Figure 5D), in *irf8-/-* retinas treated with MTZ, and in both cases decreased at 72 hpt to levels similar to the other groups (Figure 5A, C, D). The results of this analysis over time indicates that the absence of microglia delays clearance of MTZ-targeted dying rods (as expected). However, given the trajectory from 24 to 72 hpt, the increased numbers of TUNEL+ cells present in the INL and GCL of *irf8*-/- retinas at 48 hours post-Mtz treatment could represent delayed clearance of pre-existing apoptotic neurons in retinas experiencing rod damage. Alternatively, or in parallel, off-target cell death may be increased in microglia-deficient retinas upon rod ablation.

**Figure 5:**
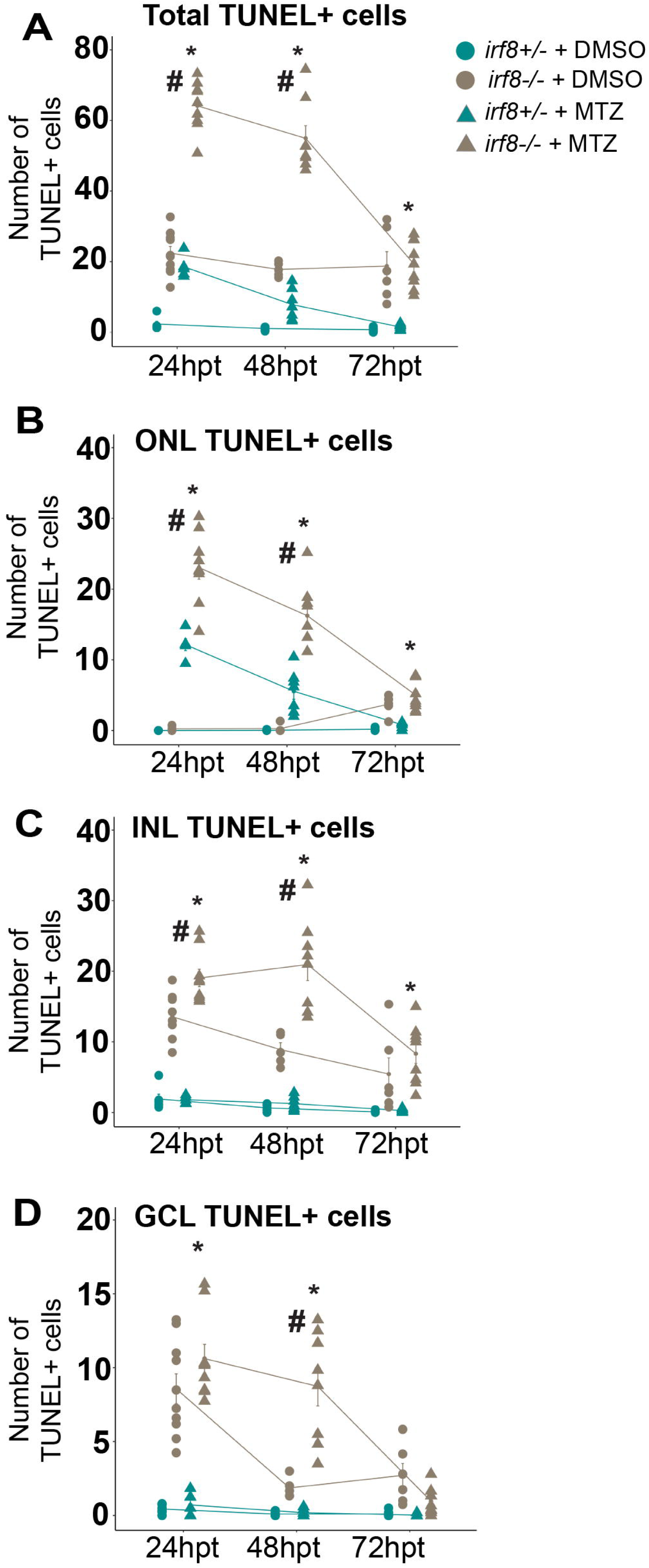
TUNEL+ cells detected following rod ablation are cleared by 72hpt in both microglia-sufficient and microglia-deficient retinas. A-D: The total number of TUNEL+ cells were counted per retinal cryosection in *rho:nfsb-eGFP* larvae of the indicated genotype and treatment. Plots show the total number of TUNEL+ cells (**A**), and TUNEL+ cells by retinal layer (**B-D**) across 24-hour intervals from 24 hpt to 72 hpt. **#** Indicates statistical significance between the groups MTZ *irf8-/-* and DMSO *irf8-/-* with p<0.05. *\** Indicates statistical significance between the groups MTZ *irf8-/-* and MTZ *irf8+/-* with p<0.05; two-way ANOVA with Tukey’s HSD post-hoc test for multiple comparisons. For clarity, only statistically significant differences are annotated in these graphs.

### Glutathione supplementation attenuates the number of TUNEL+ cells detected in the inner retina of *irf8*-/- mutants following rod ablation

Given that the detection of increased numbers of TUNEL+ cells in off-target regions upon MTZ-treatment of *rho*:nfsb-eGFP*;irf8-/-* larvae (Figures 4 and 5) is accompanied by widespread ROS (Figure 3), we hypothesized that providing antioxidant capacity to microglia-deficient retinas could potentially rescue the levels off-target cell death. We used glutathione (GSH) to supplement antioxidant capacity during treatment of *rho*:nfsb-eGFP*;irf8-/-* larvae.

Glutathione was selected because of its capacity to scavenge ROS in addition to providing reducing power for many ROS-neutralizing reactions [52,57]. We provided GSH from 24 to 48 hpt because it was in this time window that the numbers of TUNEL+ cells observed in the inner retinal layers of the MTZ-treated *irf8-/-* larvae remained elevated while the number in the DMSO-treated *irf8*-/- larvae declined (Figure 5C,D). 50µM GSH was previously shown to attenuate the detrimental effects of oxidative stress in the zebrafish retina [58]. Thus, DMSO- and MTZ treated *rho*:nfsb-eGFP*;irf8-/-* larvae were either given no GSH or a 50 µM GSH supplementation from 24-48 hpt of DMSO/MTZ treatment.

We performed these experiments in *rho*:nfsb-eGFP;*irf8*-/- retinas because the supplementation aimed to determine antioxidant rescue related to off-target death detected along with MTZ-mediated rod ablation in the absence of microglia. Treatment with GSH did not significantly impact the number of TUNEL+ cells detected in DMSO-treated *irf8*-/- retinas (Figure 6A, B, E). Consistent with previous results (Figure 4), TUNEL+ cell numbers were increased in all retinal layers of *irf8-/-* eyes upon MTZ treatment and included HuC/D+ inner retinal neurons (Figure 6C-I). Supplementation with GSH reduced the numbers of TUNEL+ cells in all retinal layers in samples receiving MTZ treatment, and this was statistically significant for the overall numbers of TUNEL+ cells (Figure 6E), TUNEL+ cells in the ONL (Figure 6F), and for TUNEL+ HuC/D+ neurons (Figure 6I). Thus, GSH supplementation provided a rescue of TUNEL+ cells detected in post-rod ablation microglia-deficient retinas. Interestingly, GSH supplementation also reduced TUNEL+ cell numbers in the ONL (Figure 6D) which at first would seem to suggest that nfsB-expressing rods may be rescued from MTZ-mediated effects. However, the number of intact, viable rod cells in MTZ treated retinas was not increased with GSH supplementation (Figure 6D, H). The reduction in TUNEL+ cells in the ONL upon GSH treatment instead may indicate that GSH supplementation impacts clearance, rather than viability, of rods exposed to MTZ in the absence of microglia. Further, the rescue of HuC/D+ TUNEL+ cell detection with GSH treatment suggests that the increased detection of these off-target retinal neurons in microglia-deficient retinas is related to changes in retinal redox biology as a consequence of microglia deficiency upon MTZ-mediated rod ablation.

**Figure 6:**
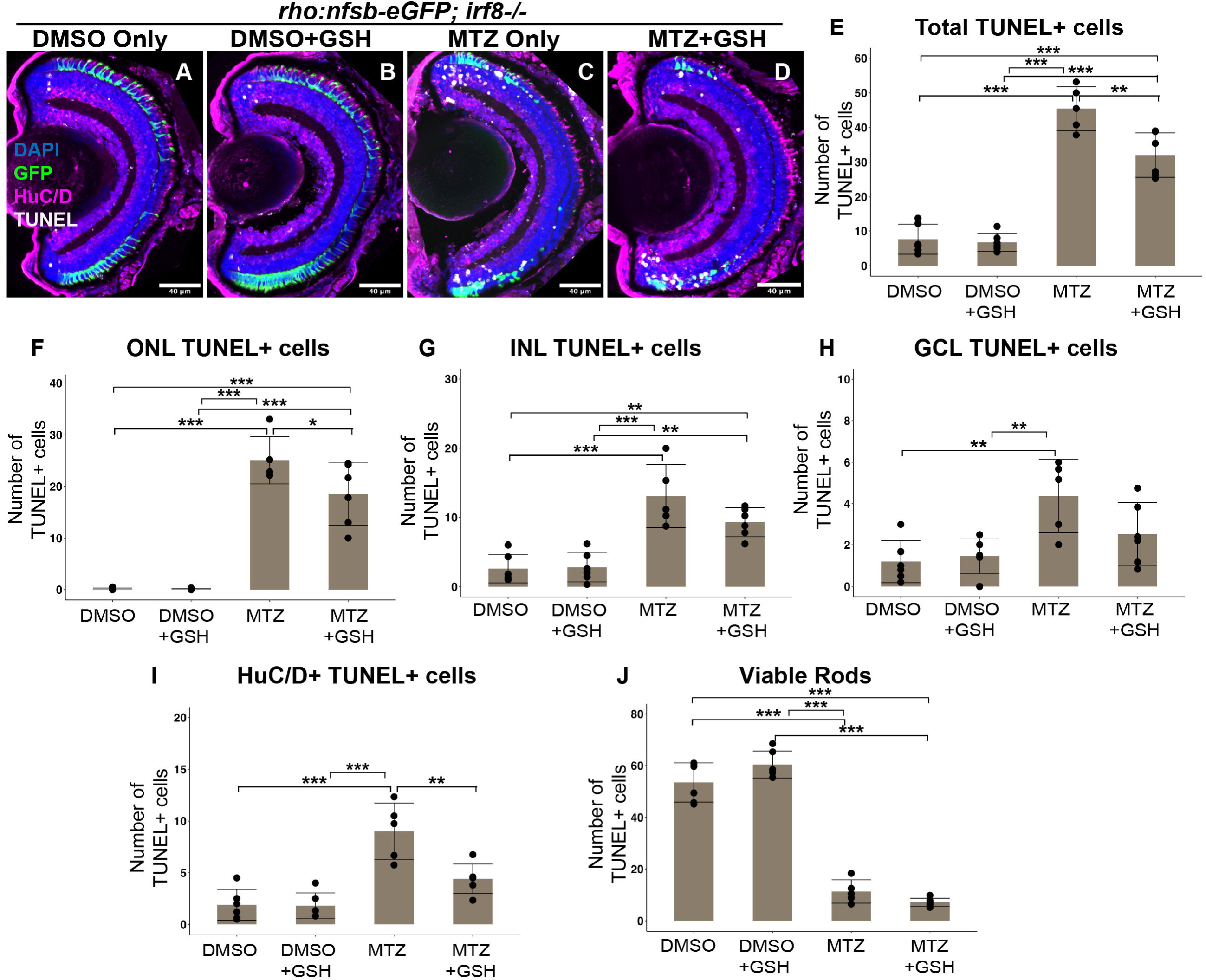
GSH supplementation rescues the number of off-target HuC/D+ TUNEL+ cells following rod ablation in microglia-deficient retinas. A-D. Representative images of retinal cryosections showing TUNEL+ cells (white), rod cells (GFP+, green), HuC/D neurons (magenta), and DAPI (blue) from *rho*:nfsb-eGFP;*irf8-/-* larvae treated with either DMSO or MTZ and receiving no supplementation (DMSO Only, MTZ Only) or supplemented with 50 µM GSH (+GSH) from 24-48 hpt. E-I: Quantifications of the total number of TUNEL+ cells (E) per retinal cryosection, TUNEL+ cells by retinal layer (F-H), and TUNEL+ HuC/D+ cells (I). (J) Quantification of the number of intact, viable rod cells per retinal section. *p<0.05, **p<0.01, ***p<0.001; two-way ANOVA with Tukey’s HSD post-hoc test for multiple comparisons. For clarity, only statistically significant differences are annotated in these graphs.

### Gene expression changes in larval eyes suggest oxidative stress and/or redox imbalance, but not glial reactivity, increase in microglia deficient retinas upon rod ablation

To further probe impacts on the retina following rod ablation in the absence of microglial responses we performed RT-qPCR to measure the expression of several genes associated with oxidative stress response (*nfe2l2a*, *txn*, *gpx1a*, *gsr*, *p62/sqstm1*) at 48 hpt in the presence or absence of rod ablation (DMSO or MTZ treatment of *rho*:nfsb-eGFP fish) in the two *irf8* genotypes (Fig 7A-E). mRNA encoding the transcription factor Nrf2 (*nfe2l2a*) was increased in *irf8*-/- compared to *irf8*+/- even following DMSO treatment (no rod ablation, Figure 7A). There was a similar trend, though not statistically significant, for expression of *txn*, *gpx1a*, and *p62* in *irf8*-/- compared to *irf8*+/- with DMSO treatment only (Figure 7B-E). Collectively, these findings suggest that oxidative stress in the developing retina is increased in the absence of microglia alone. Interestingly, mRNA levels for each of these oxidative stress genes were not significantly elevated in *irf8+/-* eyes following rod ablation (MTZ treatment, Figure 7A-E), suggesting that microglia manage ROS efficiently following MTZ-induced rod ablation. However, the levels of *nfe2l2a*, thioredoxin (*txn*) and *p62* were further elevated in *irf8*-/- eyes following rod ablation (MTZ) compared to undamaged (DSMO) *irf8-/-* eyes (Figure 7B,E). Changes in *gpx1a*, though not always statistically significant, also showed trends of increased expression in *irf8*-/- eyes following MTZ-mediated rod ablation (Figure 7C), though expression of *gsr* was not dramatically changed in these experiments (Figure 7D). It is worth noting that these experiments measure mRNA changes from bulk whole larval eyes and do not examine microglia-specific gene expression. Collectively, these results indicate oxidative stress/redox imbalance in microglia-deficient developing retinas, which is further increased in the absence of microglia upon rod ablation. These gene expression patterns are consistent with the increased detection of ROS via CellROX^TM^ upon rod ablation in *irf8*-/- retinas (Figure 3).

**Figure 7:**
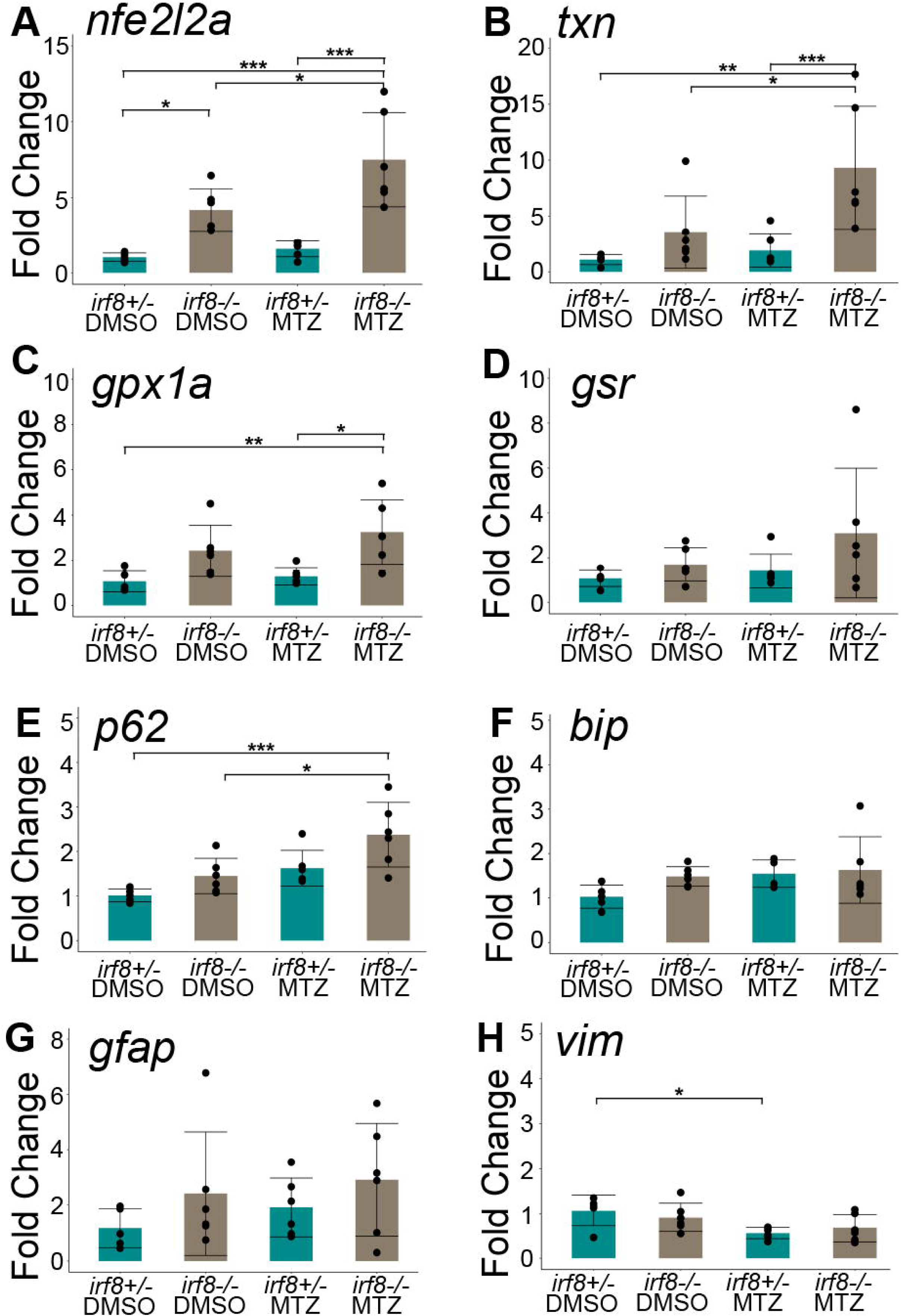
Gene expression changes in *rho*:nfsb-eGFP larval eyes measured by RT-qPCR at 48 hpt suggest increased oxidative stress and/or redox imbalance in microglia-deficient retinas. A-H: Fold change in gene expression measured by RT-qPCR using RNA/cDNA from whole *rho*:nfsb-eGFP larval eyes from the indicated *irf8* genotypes, treated with DMSO or MTZ for 48 hpt. * p<0.05, ** p<0.01, ***p<0.001; two-way ANOVA with Tukey’s HSD post-hoc test for multiple comparisons. For clarity, only statistically significant differences are annotated in these graphs.

Since oxidative stress may also be linked to ER stress, we examined expression of the endoplasmic reticulum (ER) stress-related marker *bip*, but we did not detect any changes in expression at 48 hpt in either condition or genotype (Figure 7F). Given that Müller glia respond to retinal stress and damage by increasing reactivity, which can affect the outcomes following retinal damage [59–63], we performed RT-qPCR to detect changes in the expression of genes associated with Müller glia (MG) reactivity (*gfap, vim*). Somewhat surprisingly, we did not find significant increase in the levels of transcripts for *gfap* or *vim* in response to MTZ treatment/rod ablation in *rho*:nfsb-eGFP*;irf8+/-* larvae (Figure 7G, H). These results indicate that rod ablation in larval *rho*:nfsb-eGFP retinas has limited impact on MG reactivity, which we consider to likely be a factor of age or development. Further, microglia deficiency did not result in increased levels of *gfap* or *vim* transcripts at 48 hpt in either DMSO or MTZ treated groups (Figure 7G, H), suggesting that oxidative stress or redox imbalance is not always linked to glial reactivity.

## Discussion

In this paper, we examined the consequences of microglia deficiency in the zebrafish retina following rod photoreceptor ablation using the MTZ/NTR system, with a focus on early timepoints following MTZ-induced rod ablation, primarily the first 48 hours post-treatment. The reduction of MTZ by NTR enzymes is suggested to generate ROS [43,44], and our results were consistent with this idea, as we visualized ROS in nfsB-expressing rods upon MTZ treatment using CellRox^TM^ Deep Red. We found that microglia displayed compartments filled with ROS probe signal coincident with engulfment of dying rods, indicating that they are directly challenged with this oxidative load and must manage ROS to restore homeostasis. Microglia with ROS-containing compartments were particularly detectable in the ventral retina, where rods are densely populated [50]. Despite such abundant ROS detection in dying rods and microglia, oxidative stress (as measured by gene expression of redox related genes) was minimal in the presence of microglia. Strikingly, in the microglia-deficient *irf8*-/- retinas, ROS-laden rods remained uncleared for at least 24-48 hours after their ablation and ROS signal became widespread throughout the retina in this period. This was accompanied, in microglia-deficient retinas, by increased transcript levels for genes involved in the oxidative stress/redox balance response and an increased detection of TUNEL+ cells in retinal layers outside of the ONL, including increased numbers of HuC/D+ neurons in the inner retina (“off-target” cell death).

Supplementation with glutathione (GSH) reduced the number of off-target TUNEL+ retinal neurons detected in microglia-deficient retinas following rod ablation. Collectively, our work indicates that microglial antioxidant functions are required for restoring homeostasis after MTZ-mediated rod ablation.

The increased detection of off-target TUNEL+ cells (non-rods) was a significant finding in microglia-deficient retinas following MTZ-mediated rod ablation. We noted that many of them were localized to peripheral retina adjacent to the ciliary marginal zone (CMZ), a region of stem cells and progenitors that facilitates the growth of the fish retina beyond the initial differentiation of neuroepithelium [64,65]. The location of these off-target neurons suggests that newly generated neurons may be more susceptible to changes in retinal redox homeostasis leading to their death in the absence of microglial function. It has been shown that neuronal maturation in the brain involves an attenuation of NRF2 activity [66], so it is also possible that during development of retinal neurons their antioxidant capacity may be reduced. Further, it has been reported that neurons across many conditions have limited NRF2 levels and activity [24].

Alternatively, or at the same time, these TUNEL+ neurons could represent newly generated neurons that execute programmed cell death and in normal conditions, microglia would rapidly recognize and clear them. Hence, they may be more readily detected in microglia-deficient samples, but this would also require that a compensatory mechanism for their clearance was impacted by effects of MTZ-mediated rod death. For example, this could involve the Müller glia [46], which in the presence of rod ablation in microglia-deficient retinas could be functioning to clear dying rods and could not as readily participate in inner retinal clearance. Our results are potentially consistent with both scenarios. The level of inner-retinal (INL and GCL) cell death detected at 24 hours post-treatment was similar in *irf8*-/- retinas both with and without rod ablation. At 48 hours, detection of off-target cell death (i.e., HuC/D+ neurons) remained high in microglia-deficient samples that experienced rod ablation but decreased in microglia-deficient samples that did not. Without live timelapse imaging, it is extremely challenging to determine whether the increased number of TUNEL+ neurons in microglia deficient retinas experiencing rod ablation is due to off-target cell death, delayed compensatory clearance, or a combination of both. While we were able to image whole, dissected eyes following fixation and methanol-mediated clarification of pigment at the ages involved in these experiments, we faced limitations to live imaging due to pigmentation that, over time, becomes PTU-insensitive, increased eye size due to growth, and the cumbersome number of transgenic reporters that this would require. Nonetheless, off-target cell death detection was partially rescued by GSH antioxidant supplementation, suggesting that the antioxidant functions of microglia may underlie the restoration of homeostasis.

Regardless of the origin of the increased numbers of TUNEL+ cells at 48 hours post rod ablation in microglia-deficient retinas, our results indicated that clearance of these “off-target” TUNEL+ cells and the on-target ablated rods occurred by 72 hours. We posit that the retinal Müller glia are the most likely candidate for such compensatory clearance. We have shown that in the absence of microglia, Müller glia increase engulfment of dying neurons in the developing zebrafish retina [46]. More recently, using in vivo real-time imaging we directly documented phagocytosis of apoptotic neurons and lysosomal fusion with cargo containing compartments in Müller glia [1] attesting to their ability to perform this function. However, phagocytosis and cargo digestion by Müller glia took comparatively longer than for microglia, and microglia were clearly the primary phagocyte in developing zebrafish retina [1,2]. Therefore, it follows that microglia are likely suited to perform rapid antioxidant function when faced with cargo containing ROS or other oxidants.

In line with the idea of increased oxidative stress on the retina in the absence of microglia, we observed that some of the diffuse ROS signal in microglia-deficient retinas localized to Müller glia cells. This could indicate that Müller glia compensate for oxidative load management in the absence of microglia, or that secondary ROS are generated and/or propagated due to delayed sequestration of ROS via engulfment of dying rods when microglia are absent. Surprisingly, glial reactivity (as reported by *gfap* and *vimentin* mRNA levels), and the ER stress gene *bip*, were not increased in *irf8*-/- retinas in the presence or absence of rod ablation. This does not rule out the possibility of ER-stress, but we did not detect any indication through the expression of this particular marker. However, several oxidative stress genes showed increased mRNA levels in microglia-deficient retinas that were further elevated upon rod ablation. Together, these results could represent a compensatory antioxidant response by the Müller glia, or even other retinal cells such as neurons and the retinal pigment epithelium (RPE), aimed to restore homeostasis in the absence of microglial antioxidant function. Further, our results indicate that oxidative stress is not always linked to Müller glia reactivity.

We observed apparent effects of MTZ in non-transgenic (no nfsB/NTR expression) *irf8* mutants, namely the detection of CellROX^TM^ signal and TUNEL+ cells, both observed in the inner retina. This suggests that microglia could be important for detoxification of the pro-drug in the absence of the nfsB reaction. The pro-drug itself may generate a low level of superoxide in the presence of oxygen in a futile-redox cycling process [43] and/or have other consequential effects that microglia contain effectively. Such detoxification activity may utilize enzymes readily expressed by microglia, and in their absence, this burden may be shifted to other retinal cells such as the Müller glia. Nonetheless, impacts of microglia deficiency on detection of reactive oxygen species and cell death markers were markedly more pronounced when nfsB-expressing rods were ablated upon MTZ treatment.

Microglia are well known for their clearance functions in response to neuronal death, but how they mitigate high oxidative loads obtained through phagocytosis remains to be determined. Our previous work [31] and results here support that Nrf2 and some of its known transcriptional targets are increased at the level of mRNA expression, but it is not clear which antioxidant gene products are most crucial. Numerous enzymes are involved in complex biochemical reactions that neutralize ROS [52,57,67]. Further, CellROX^TM^ has been described to principally detect superoxide and hydroxyl radicals [47,48], but this does not exclude the generation of other free radicals and reactive nitrogen species (RNS) via the MTZ/nfsB reaction that we did not visualize/detect. The exact mechanism of ROS generation by the transgene-encoded nfsB enzyme in eukaryotes has not been specifically established to our knowledge, although there are insights that propose a mechanism by which ROS arise following the nitroreductase reaction [43,44]. Yet to be further investigated is the impact on regenerative responses by the Müller glia, which are also the source of regenerated retinal neurons in zebrafish [68–74] and as discussed above, are likely impacted by oxidative stress in microglia deficient retinas. Several papers indicate microglia influence retinal regeneration in zebrafish [40,54,75–78], but this has not yet been connected to their putative antioxidant functions.

Early responses of microglia are important to understand in order to mitigate pathology following acute trauma but may also provide insight to compounding or accumulating molecular and cellular changes that occur during neurodegenerative disease. Our work here emphasizes that microglia, through antioxidant functions, contribute to a timely restoration of homeostasis following the death of photoreceptors that is intertwined with ROS/oxidative stress. ROS (or other free radicals) may accumulate over time in neurodegeneration and alongside microglial dysfunction, could lead to secondary pathology. There is likely a link between redox state, reactive oxygen species (ROS), and the inflammatory state and signaling of microglia such that deficient activity of the antioxidant pathway contributes to progression of pathology [79–82]. The opposite has also been documented, wherein augmentation of the antioxidant pathway is beneficial for neuroprotection and injury repair [83–86]. Additionally, our work suggests that compensatory function of other glial cells may be executed to mitigate oxidative stress and restore redox balance when microglia are unable to perform such functions. Our findings here show that oxidative stress levels may be elevated in developing retinas when microglia are absent even when additional insult is not applied, and this may indicate additional load or function required from the retinal Müller glia. Such compensatory activities of glia have recently been recognized by our lab and others [46,87–89], and likely represent contexts relevant to neurodegenerative disease.

## Methods

### Animals

All procedures were in compliance with protocols approved by the Institutional Animal Care and Use Committee at the University of Idaho. Adult zebrafish (*Danio rerio*) were housed in an aquatic vivarium with filtered, recirculating water, maintained at 28.5°C, and an automated 14h light:10 h dark cycle following procedures outlined in *The Zebrafish Book* [90]. Zebrafish embryos were collected into glass beakers from spawn tanks in the morning with light onset considered to be 0 hours post-fertilization. Zebrafish cannot be sexed before reproductive maturity, therefore sex of larval zebrafish used in the experiments could not be determined.

Embryos that were used for whole eye imaging were housed in an incubator, with 14-10h light-dark cycle, at 28°C and treated with N-Phenylthiourea (PTU) at a final concentration of 0.006% in order to prevent the development of pigmentation. The water and PTU were refreshed each day. Prior to collection of larvae at experimental endpoints, larvae were terminally anesthetized in a working solution of 0.25 mg/mL MS-222/tricaine (Syndel USA).

### Zebrafish lines

The transgenic line *rho*:nfsb-eGFP*^nt19* was generated and gifted by Dr. David Hyde’s lab (University of Notre Dame) [42]. In this line the *rho* promoter drives the expression of nfsb-eGFP fused to eGFP fluorescent protein. Upon receipt in our facility, the *nt19* line was outcrossed to our wild-type strain referred to as SciH, originally obtained from Scientific Hatcheries (now Aquatica Tropicals), and to *irf8*^st95 carriers. The *irf8^st95* line was generated and gifted previously by Dr. Celia Shiau (UNC) [45]. Genetic crosses were performed to produce fish carrying the *rho*:nfsb-eGFP transgene and heterozygous or homozygous for the *irf8*^*st95* allele; these are referred to throughout the manuscript as *irf8+/-* and *irf8-/-*, respectively. Expression of the *rho*:nfsb-eGFP transgene was determined by viewing embryos at 3 days post-fertilization (dpf) through a fluorescent sorting microscope and visibly scoring GFP expression in eyes.

Experiments were performed with clutches of both *irf8* genotypes, and genotype unknown at the time of the experimental procedures. Genotype of the larvae was determined post-fixation from tail clips using a PCR/Restriction digest assay as described in [45].

### Rod ablation

Treatment of larval zebrafish began at the age of 3.5 dpf. Larvae were immersed in a solution of 10mM metronidazole (MTZ) in 0.1% by volume dimethyl sulfoxide (DMSO), or in a solution of 0.1% DMSO (vehicle). Solutions were prepared in fish system water and protected from light during mixing. Treatments were performed in a Heratherm™ incubator set at 28°C. Because MTZ is light-sensitive, larvae exposed to DMSO/MTZ solutions were treated in petri dishes and protected from light by loosely covering dishes with aluminum foil. After 24 hours of treatment, larvae received a fresh solution of their respective treatment and then returned to the incubator for another 24 hours. Following the duration of treatment (48 hours post treatment, hpt), larvae were removed from dishes and either fixed for tissue preparations (described below) or transferred to a pulse of CellRox™ immersion. For samples collected at 72 hpt, larvae were transferred to clean water for recovery and collected 24 hours later.

### CellRox™ immersion

At the end of the DMSO/MTZ treatment, larvae were removed from treatment solutions and placed in a solution consisting of 0.1% CellRox™ Deep Red reagent (Invitrogen, working concentration of 2.5 µM) and 0.006% N-phenylthiourea in fish system water. Larvae were incubated in CellRox™ solutions for 30 minutes at 28°C and protected from light. Following incubation, larvae were removed from solution, terminally anesthetized in a working solution of 0.024% MS222-tricaine, then fixed in a solution of 4% paraformaldehyde in 1x PBS.

### Immunofluorescence of whole tissue

At designated experimental endpoints, larvae were fixed in a solution of 4% paraformaldehyde in 1x phosphate buffered saline (PBS) overnight at 4°C. The next day, the fixation solution was removed, and larvae were rinsed once in 1X PBS+ 1% Triton-X100 (PBST) then three times in 1X PBS for 5 minutes each. Following PBS rinses, the larvae were dehydrated with four x 100% methanol washes for 10 minutes each, then stored in 100% at −20°C. Prior to staining, methanol was removed from the samples, and they were rehydrated with two 15-minute washes of 1x PBST (1% triton-X-100). Following rehydration, the blocking step was performed by incubation in antibody dilution buffer (1% normal donkey serum, 0.1% of sodium azide, 0.1% triton-X-100 in 1X PBS) for at least one hour at room temperature. The primary antibody solution was prepared in antibody dilution buffer. Primary antibodies were used as follows: mouse anti-4C4 (1:100, hybridoma supernatant from clone 7.4.C4), mouse anti-glutamine synthetase (1:100, BD Biosciences). Samples with primary antibody mixtures were incubated overnight at 4°C with constant rocking. Following primary antibody incubation, the samples were washed with 500 µL of 1x PBST (1% triton-X-100) for 30 minutes at room temperature. Secondary antibody solutions were prepared at 1:200 dilution in antibody dilution buffer. Secondary antibodies included goat anti-mouse Cy7 (1:200, AAT Bioquest). DAPI was included in the secondary antibody solution at a dilution of 1:1000. Samples were incubated in secondary antibody solution overnight at 4°C with constant rocking. The next day, the samples were washed again with 500µL of 1x PBST for 30 minutes at room temperature, followed by a 20-minute rinse with 1x PBS.

For imaging, eyes were dissected from fixed and stained whole larvae and suspended in glycerol. A “coverslip sandwich” was made using a 22×60 mm microscope slide coverslip and attaching two smaller 22.5×22.5 mm coverslips on each end of the longer coverslip using adhesive. Samples were mounted in the space on the 22×60 mm coverslip between the 22.5×22.5 mm coverslips. When the samples were oriented, a second 22×60 mm coverslip was attached by placing on top of the smaller coverslips to create a “sandwich” effect.

### Fluorescence In-Situ Hybridization

Fluorescence in-situ hybridization (FISH) was performed using hybridization chain reaction (HCR) reagents and protocol from Molecular Instruments [91] as done previously [1,53]. The NCBI accession numbers of the genes of interest (**Table 1**) were provided to Molecular Instruments for the design and synthesis of gene-specific probe sets. At the experimental endpoint, larvae were fixed in a freshly made solution of 4% paraformaldehyde in 1x phosphate buffered saline (PBS). The PFA solution was cooled at 4°C prior to use for fixation. The samples were fixed in 1mL of fixation solution overnight at 4°C on a nutating mixer. Following fixation, the PFA solution was removed, and samples were washed three times with 1mL RNAse-free 1x PBS for five minutes per wash. The samples were then washed four times with 1mL of 100% methanol for 10 minutes per wash. Afterwards, the samples were stored in 1 mL of fresh 100% methanol and stored at −20°C at least overnight.

**Table 1.**

| Gene | Accession number | Lot number | Hairpin/amplifier |
| --- | --- | --- | --- |
| *mpeg1* | NM_212737.1 | PRP656 | B3-Cy7 |
| *gpx1a* | NM_001007281.2 | PRO917 | B2-AF647 |
| *gsr* | NM_001020554.1 | PRO918 | B1-AF647 |

To begin the HCR-FISH procedure, samples were first rehydrated in a graded series of methanol-1x PBS-Tween (0.1%) solutions for 5 minutes per rehydration with 25:75, 50:50, and 75:25% ratios of PBS-Tween:MeOH before a final step of 100% 1x PBS-Tween. Samples were treated with 1mL of 10µg/mL Proteinase K solution for 10 minutes at 37°C. Following Proteinase K treatment, the samples were briefly rinsed with 1x PBST and then post-fixed with a solution of 4% PFA in 1x PBS for 20 minutes at room temperature. Following the post-fix step, the samples were once again washed thrice with 1x PBST for 5 minutes per wash. Following the PBST washes, the samples then underwent a pre-hybridization step in which they were incubated in probe hybridization buffer pre-heated to 37°C; this step was carried out at 37°C in a hybridization oven for 30 minutes. Afterwards, the probe hybridization solution was prepared for the samples by combining 500 µl of probe hybridization buffer with 2 pmol of each desired probe set in accordance with the appropriate multiplexing considerations. This ratio was followed for each tube of samples. Samples were incubated with the probe set solution for 12-16 hours at 37°C in a hybridization oven.

After hybridization, the probe solution was removed, and samples were washed with a probe wash buffer four times for 15 minutes per wash at 37°C followed by 2 washes in a 5x saline sodium citrate-0.1% Tween-20 buffer (5xSSCT) for 5 minutes per wash. Following the washes, samples were incubated in probe amplification buffer for 30 minutes at room temperature. An amplification solution was then prepared (500µl/tube) with hairpin amplifiers. First, 30 pmol of hairpins (per tube of reaction) were snap cooled by heating the hairpin at 95°C for 90 seconds in a thermocycler, followed by cooling to room temperature in the dark for 30 minutes. Once the hairpins had cooled, they were added to 500µl of room temperature amplification buffer in accordance with the appropriate multiplexing considerations. Samples were incubated in the hairpin mixture for 12-16 h at room temperature, protected from light.

Following amplification, the hairpin solution was removed, and samples were washed with SSCT five times. DAPI was added at 1:1000 in the final wash, and then samples were stored in 1× PBS at 4°C until imaging. Imaging was performed within 24 hours. Eyes were dissected from the body, suspended in glycerol, and mounted in “slide sandwiches” as described above.

### Preparation of retinal cryosections

At designated experimental endpoints, larvae were fixed in solution of 4% paraformaldehyde in a 5% sucrose phosphate buffer and incubated overnight at 4°C. Afterwards, the fixative was removed, and the samples were washed in 5% sucrose phosphate buffer thrice for 10 minutes each. The samples were then washed with graded series of sucrose phosphate solutions beginning with a 2:1 ratio of 5%:20% sucrose phosphate buffer, followed by 1:1, and finally 1:2 ratio for 30 minutes each. Finally, the samples were incubated with a 20% sucrose phosphate buffer overnight at 4°C. Samples were then embedded into tissue blocks composed of a medium of 2:1 20% sucrose phosphate buffer: OCT and stored at −20°C at least overnight. Tissue blocks were sectioned using a Leica CM3050 cryostat with 10µm cryosection thickness, dried in a desiccator, then stored at −20°C.

### TUNEL assay and immunostaining of retinal cryosections

Microscope slides with retinal cryosections were thawed for 10 minutes at room temperature following removal from the freezer. Because of the extant GFP fluorophore present in the tissue, the samples were protected from light as much as possible during active procedures and during all incubation steps. TUNEL staining was performed following the manufacturer’s instructions (Roche). After thawing, slides were rinsed in 1x PBS solution for 20 minutes. During this incubation, sodium citrate permeabilization solution was prepared on ice following the manufacturer’s instructions. Slides were then placed in the cooled permeabilization solution for 2 minutes on ice then immediately removed and placed into 1x PBS for a 5-minute rinse on an orbital shaker. This rinse step was repeated once. During the rinse, the TUNEL reaction mixture (TUNEL TMR Red, Roche, Cat. 12156792910) was prepared on ice according to the manufacturer’s instructions. Excess PBS was removed by gentle wicking. Then a liquid blocker gel pen was used to draw a circle around the tissue in order to provide a barrier for the small volume of TUNEL reaction mixture to stay in place. 15µL of TUNEL reaction mixture was placed on each section. The slides were then placed in an Broekel Scientific™ InSlide Out™ hybridization oven on a slide tray and incubated for 3 hours at 37°C. For humidification, pieces of filter paper were partitioned and placed in the slide tray and overlayed with approximately 5mL of distilled water. Following the reaction incubation, the slides were rinsed with three 1x PBS rinses of 5 minutes each.

Immunostaining immediately followed the TUNEL procedure. Tissue was blocked for at least 1 hour at room temperature in blocking solution (described above), then incubated with primary antibody solution at 4°C overnight. Primary antibody solution was prepared in antibody dilution buffer with rabbit anti-HuC/D (1:200, Abcam ab210554). The next day, slides were washed in 1x PBST (1% Triton-X-100) for 30 minutes at room temperature. The secondary antibody solution was prepared with donkey anti-rabbit AlexaFluor647 (Jackson ImmunoResearch, 1:200) and DAPI at a dilution of 1:1000. Following the PBST wash, the slides were incubated with secondary antibody solution for 2 hours at room temperature. Slides were once again washed with 1x PBST for 30 minutes at room temperature, followed by a 20-minute wash in 1x PBS. Stained tissue was mounted in Vectashield Vibrance® (Vector Labs), covered with microscope slide coverslips, and cured = in the dark at room temperature for at least two hours prior to imaging.

### Confocal Microscopy

Images were acquired on a Nikon Eclipse Ti2-E, Crest X-light V3 spinning disk confocal microscope using a CFI APO LWD 40x water immersion 1.15 NA λ S DIC N2 objective. Image acquisition was performed using a computer workstation running Nikon Elements software. For whole eyes, image stacks were acquired with 3μm z-steps. For retinal cryosections, images were acquired with 2µm z-steps. Imaging of the CellRox™ Deep Red dye was done using excitation with a far-red laser (∼640 nm). Imaging of the Cyanine 7 (Cy7) fluorophore was done with a Cy7 excitation laser line (∼711nm). DAPI was imaged using the DAPI laser line (365 nm). Imaging of GFP was performed using a GFP laser line (∼480 nm), and imaging TUNEL TMR Red was performed using a red laser line (∼560 nm).

### Image viewing and preparation

Image processing and quantifications were performed using ImageJ/FIJI. Viewing of both individual z-stacks and maximum projection images of select z-stacks was used to analyze according to the intended metric. A cell counter plug-in was used to count cells of interest (e.g., TUNEL+ cells). Microscopy images presented in our figures were converted into RGB tiff files from their original 16-bit format. Movies were made by selecting a range of z-stacks and displaying specific channels and then saving the montage as an .avi file with JPEG compression.

### Quantification of fluorescence intensity

Quantification of fluorescence intensity in confocal images was performed using FIJI/Image J. Fluorescence intensity was determined using single z-stack images and was adapted from our lab’s previous publication [46]. Prior to measurement, the measurement options in Image J were set to include area and integrated density. For CellRox™ fluorescence, the intensity of the CellRox™ signal was measured by drawing a region of interest (ROI) around a microglial cell determined by the shape of the signal of the 4C4 antibody stain. For the intensity of mRNA transcript via HCR-FISH, the bounds of the microglia were approximated by the bounds of the *mpeg1* transcript signal. The measured intensity of the target signal of interest was then measured within those bounds to serve as the approximate intensity within microglia. The approximate median z-stack of a cell was chosen for fluorescence measurement. In addition to the fluorescence intensity within the bounds of the cell, several regions of background outside the cell were measured and averaged to provide mean background signal value. From here, the average background was multiplied to the area of the region of interest to generate a value of mean background signal. That generated value was subtracted from the integrated density to yield the corrected total cell fluorescence (CTCF).

### RT-qPCR

At the experimental endpoint, anaesthetized larvae were placed in a 1.5mL Eppendorf tube, water was removed, and larvae were flash frozen on liquid nitrogen and immediately overlayed with super-chilled 100% methanol. Samples were stored at −80°C until ready for RNA extraction. For eye dissection, the samples were held on ice and eyes were removed from larval heads and placed into RNA lysis buffer (Zymo Quick-RNA microprep kit). Tissue was homogenized in lysis buffer using a handheld homogenizer (Bio-Gen Series PRO200, ProScientific). RNA was extracted Zymo Quick-RNA microprep kit (Zymo Research) according to the manufacturer’s protocol. Each sample was a pool of both eyes from a single fish. For genotyping of samples, the remainder of the body was transferred to 25µl NaOH and gDNA prepared as established in *Meeker et al.*[92]. DNA was used for genotyping as described above. Following RNA extraction, cDNA was prepared using SuperScript IV cDNA synthesis kit (Invitrogen). cDNA was stored at −20°C until use.

Quantitative PCR (qPCR) was performed using PowerTrack SYBR Green Master Mix (Applied Biosystems), with 1 ng of cDNA per reaction. Reactions were performed in duplicates. Primers for genes of interest were selected from previous publications and are listed in Table 2. The qPCR reaction was carried out in a QuantStudio™ 3 Real-Time PCR System (ThermoFisher) under standard reaction program settings with 40 amplification cycles. β-actin served as the housekeeping gene for our analysis, and fold-change was determined by the ΔΔCT method.

**Table 2:**
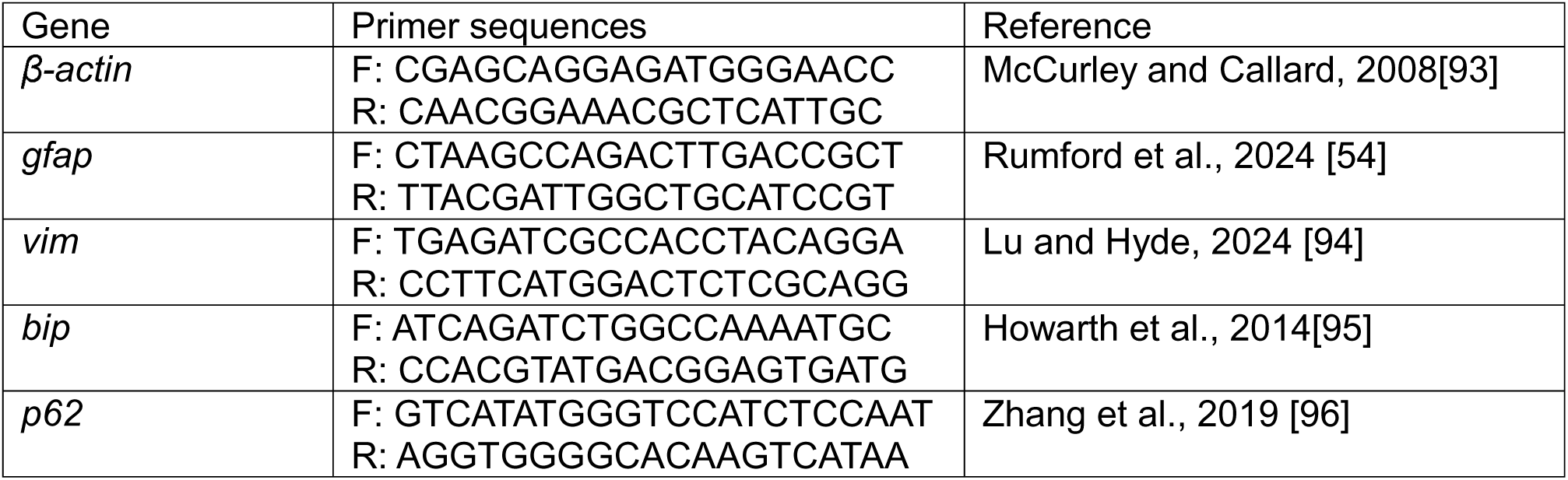
Primers used for qPCR.

### Statistical Analysis

Statistics were computed in R-Studio. For the statistical comparisons in Figures 1 and 2, a T-test was performed. For comparisons between multiple groups, a two-way ANOVA test was performed followed by Tukey’s honest significant difference post-hoc test. P-values less than 0.05 were considered statistically significant.

## Acknowledgements

We thank Dr. David Hyde for gifting the *rho*:nfsb-eGFP^nt19 transgenic line, and Dr. Celia Shiau for previously gifting the *irf8*^st95 mutant zebrafish. We thank the University of Idaho Imaging and Data Acquisition Core (IDAC, managed by the Institute of Modeling Complex Interactions/IMCI) and Institute for Interdisciplinary Data Science (IIDS) for support of our research. We thank Parker Van Dam (University of Idaho) for technical assistance with several experiments, and Jordan Rumford (University of Idaho) for assistance with fish care and husbandry. We thank Dr. Deborah Stenkamp (University of Idaho) for critical review of earlier versions of the paper.

## Author Contributions

MM performed data acquisition and data analysis. MM and DMM performed data interpretation. DMM and MM conceived the project, generated the figures, and wrote the manuscript.

**Supplemental Figure 1: CellRox™ signal in whole eyes of *irf8-/-* larvae is elevated by rod ablation**. A-A’: Images of CellRox™ signal in whole larval eyes from DMSO-treated larvae at 48 hpt. A: Image of a DMSO-treated whole eye at 48 hpt from *rho*:nfsB-eGFP- (non-transgenic) larva. A’: Image of a whole eye from *rho*:nfsB-eGFP+ larva at 48 hpt. B-B’: Images of whole larval eyes from MTZ-treated larvae at 48 hpt. B: Image of whole eye from *rho*:nfsB-eGFP- (non-transgenic) larva at 48 hpt. B’: Image of whole eye from *rho*:nfsB-eGFP+ larva treated with MTZ at 48 hpt; image was previously shown in Figure 3.

## Movie Captions

**Movie 1:** Confocal z-stack series from images take from whole larval eye of *rho*:nfsb-eGFP;*irf8*-/- fish following MTZ-induced rod cell death at 48 hpt. CellRox™ signal (magenta) and staining for glutamine synthetase (GS, to label Müller glia, cyan) are shown.

**Movie 2:** Confocal z-stack series showing ventral region of whole larval eyes from *rho*:nfsb-eGFP transgenics from *irf8*+/- (left) and *irf8*-/- (right) genotypes following MTZ-induced rod ablation at 48 hpt. Images show rods (green), microglia (yellow), CellRox™ probe signal (magenta), and cell nuclei/DAPI (blue). Pyknotic nuclei appear as condensed, bright blue nuclei. Note the absence of microglia in images from *irf8*-/- eyes.

