## Supplementary figures and images for "Microglia maintain retinal redox homeostasis following ablation of rod photoreceptors in zebrafish"

### Supplemental Figure 1

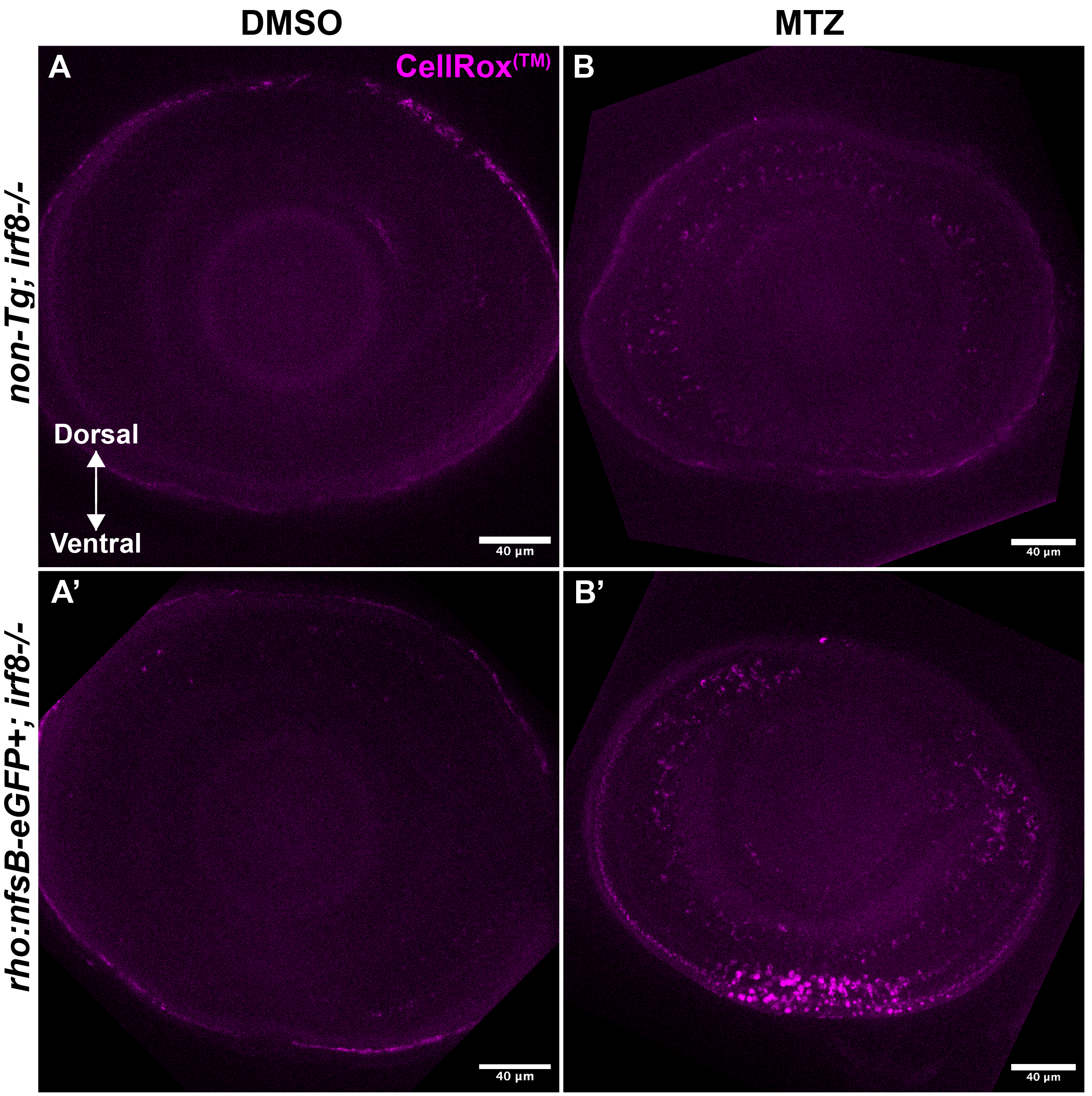
